# Cicada timetree by BEAST v1.X verified the recently and exponentially increasing base substitution rates

**DOI:** 10.1101/2020.12.03.409599

**Authors:** Soichi Osozawa, John Wakabayashi

**Author notes:** **Corresponding author.** (S. Osozawa).

## Abstract

Following the recent publication of global cicada phylogenetic trees by Marshall et al. (2018), Łukasik et al. (2018), and Simon et al. (2019), we developed a new dated tree incorporating mostly endemic east Asian cicada data for totally 113 specimens, using the mostly advanced BEAST v1.X software applied the relaxed clock model. Fossil calibrations as old as Triassic were adopted after Moulds (2018), and a Quaternary geological event calibration was adopted following Osozawa et al. (2012), applying the calibration function of BEAST. Our timetree suggests that Tettigarctidae had cicada basal lineage as old as 200 Ma, and Derotettiginae was next as old as 100 Ma. Tibicininae was a sister of the resting Cicadidae, and Tettigomyiinae, Cicadettinae, and Cicadina started simultaneous branching and radiation around 40 Ma. We made a base substitution rate vs age diagram based on the timetree using the BEAST function, and it strongly suggested an exponential increase of base substitution rate approaching the present. The consequent increased cicada biodiversity including generation of cryptic species might have been driven by the generation and spreading of C4 grasses and the following Quaternary glaciations and severe environmental change.

## Introduction

A phylogenetic tree of worldwide cicada was recently constructed by Marshall et al. (2018) and Simon et al. (2019), applying five concatenated sequences of mitochondrial COI and COII, and nuclear ARD1, EF-1a, and 18S rRNA, and by Łukasik et al. (2019), applying whole mitochondrial sequences for representative species.

These Simon group studies included comparatively little Asian data. After Osozawa et al. (2017), which treated *Platypleura* endemic cicadas in east Asia, mostly from the Japan, Ryukyu, and Taiwan islands, we collected cicada samples and increased the available DNA sequence data of mitochondrial COI and nuclear 18S rRNA (Table 1; totally 92 specimen data / 92 × 2 = 184 sequence data) were newly added to the already published 70 specimen data of *Platypleura* (duplicated sequence data were eliminated, and totally 21 specimen data were considered to construct the present timetree). Our east Asian data combined with the global data offered by Marshall et al. (2018; including 20 east Asian species / totally 149 species in their figure 4) and Łukasik et al. (2018; including 27 east Asia species / totally 93 species in their figure 2) allows the construction of a more complete phylogenetic tree, and the operational taxonomic units (OTUs) including outgroup species are 191 for the COI timetree in Figs. 1 and 2, and 155 for the COI +18S rRNA timetree in Fig. 3. Note that the COI and 18S rRNA data in Table 1 by Marshall et al. (2018) contain many missing and incompatible data, and their partial 42 data / total 149 data (our own data: 113) were only available in our analyses.

**Table 1.**
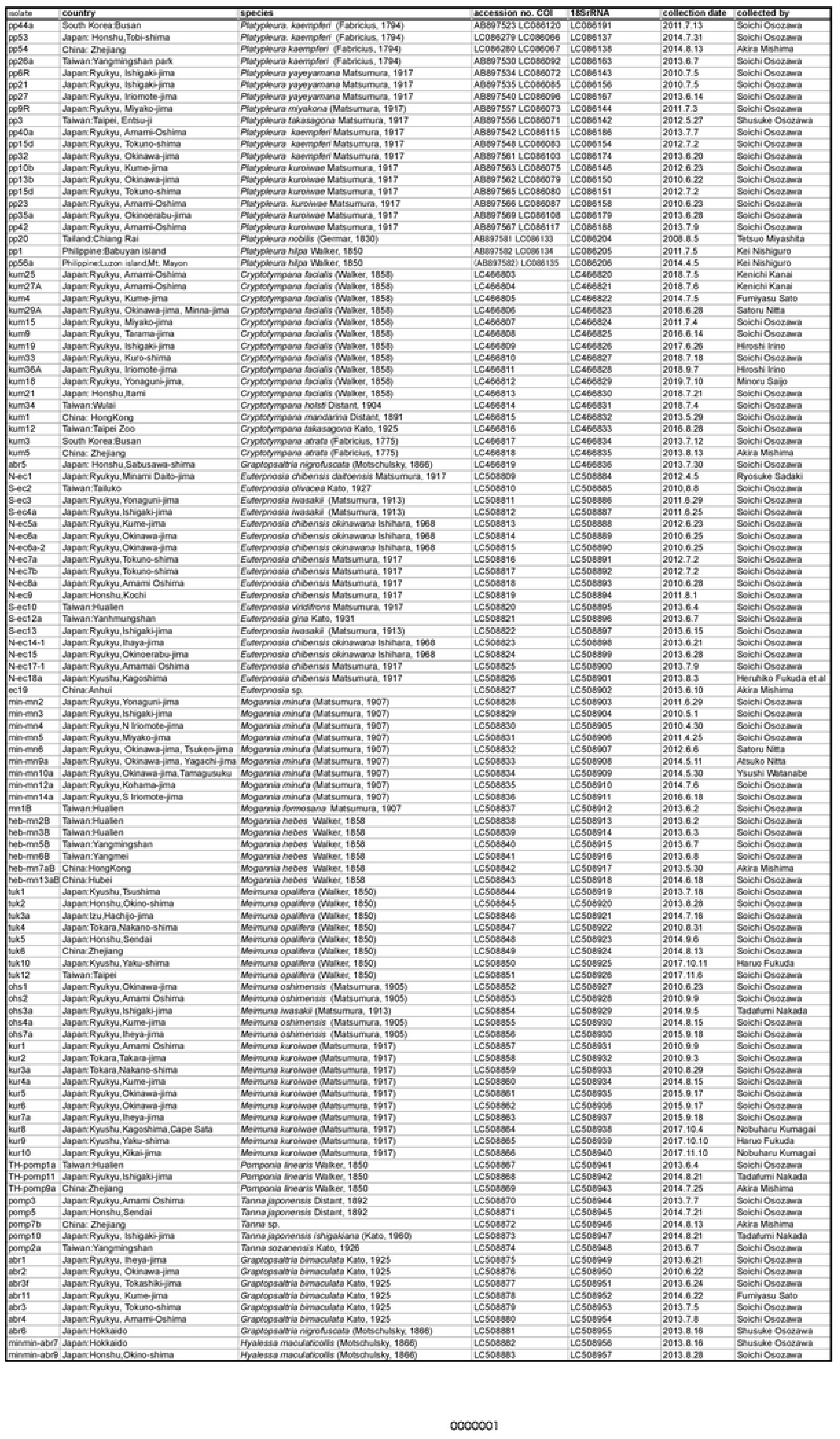
East Asian cicada species collected and analyzed in this paper.

**Fig. 1.**
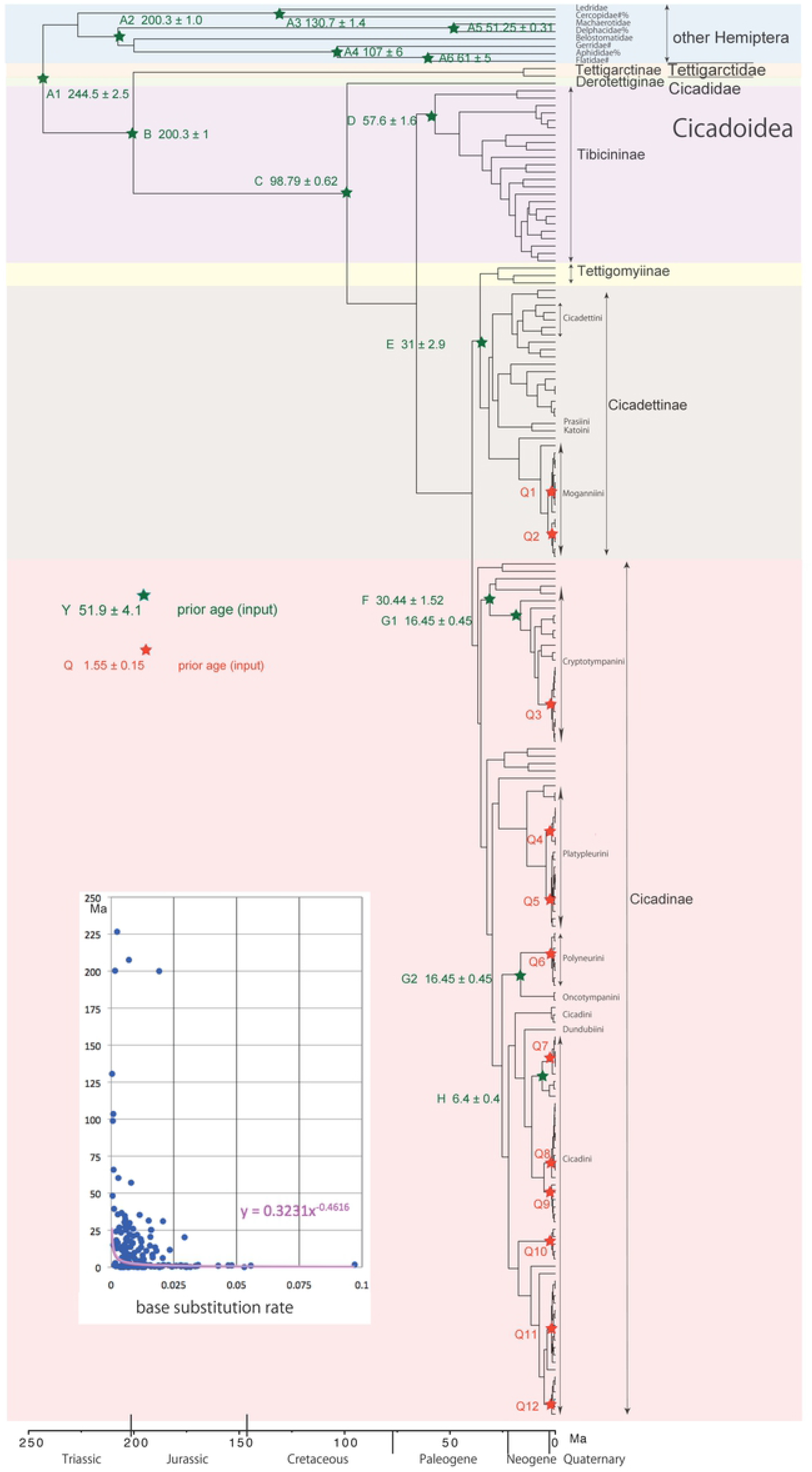
Simplified cicada timetree built by BEAST v1.X, applying 1,534 bp COI sequence. Inserted figure: Base substitution rate (= rate median shown at each node; mutations per bp per million years) vs age (= posterior age shown at each node) diagram. Purple curve with equation: Exponential trendline calculated using Excel.

**Fig. 2.**
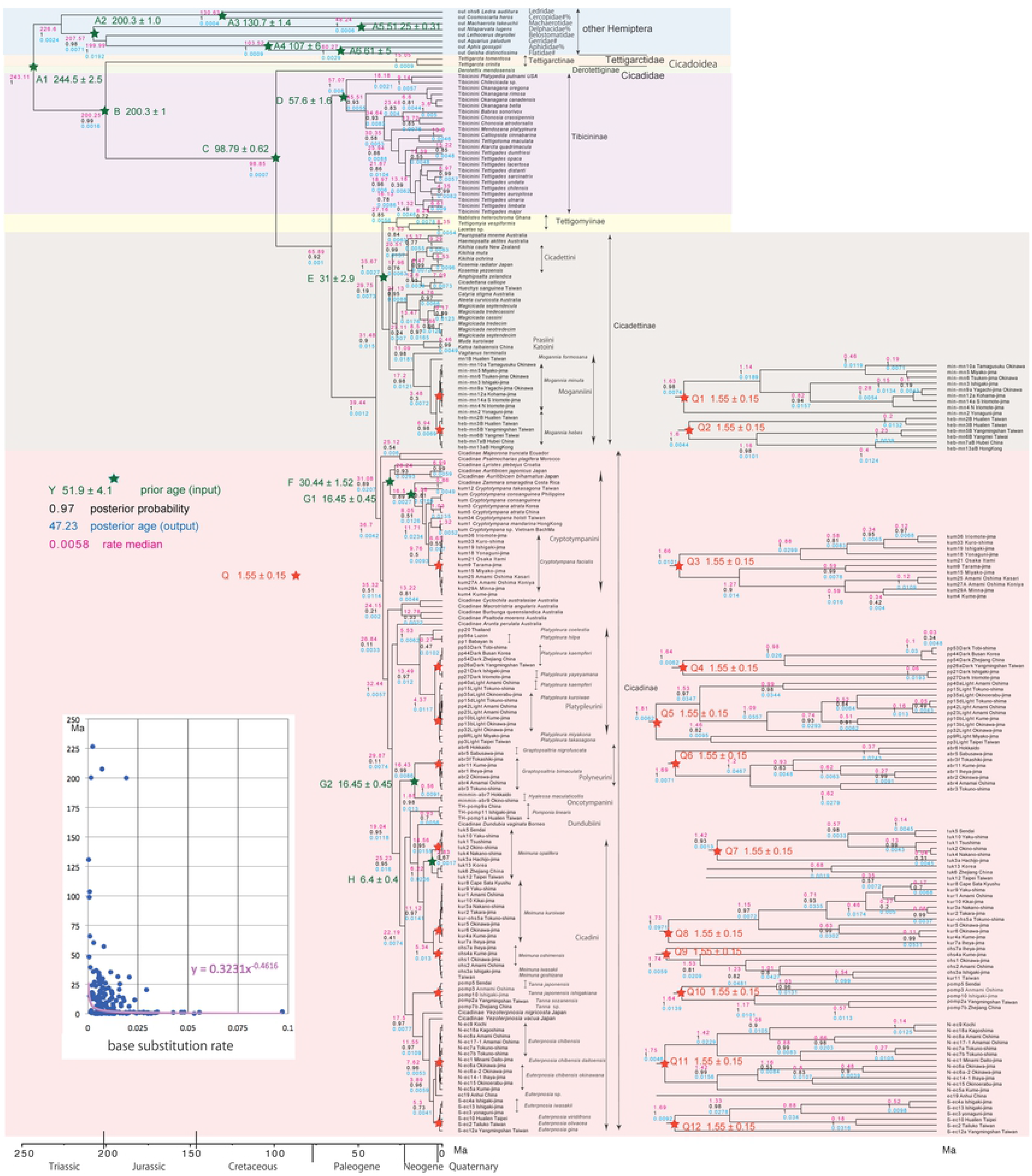
Cicada timetree built by BEAST v1.X, applying 1,534 bp COI sequence. OUTs with isolate number: our own analyzed specimens shown in Table 1, and others: from GenBank / DDJB. In outgroup Hemiptera, #: analyzed family by Johnson et al. (2018), % analyzed family by Misof et al. (2014). Note that the outgroup is calibrated by the single point point A1. Inserted figure: Base substitution rate (= rate median shown at each node; mutations per bp per million years) vs age (= posterior age shown at each node) diagram. Purple curve with equation: Exponential trendline calculated using Excel.

**Fig. 3.**
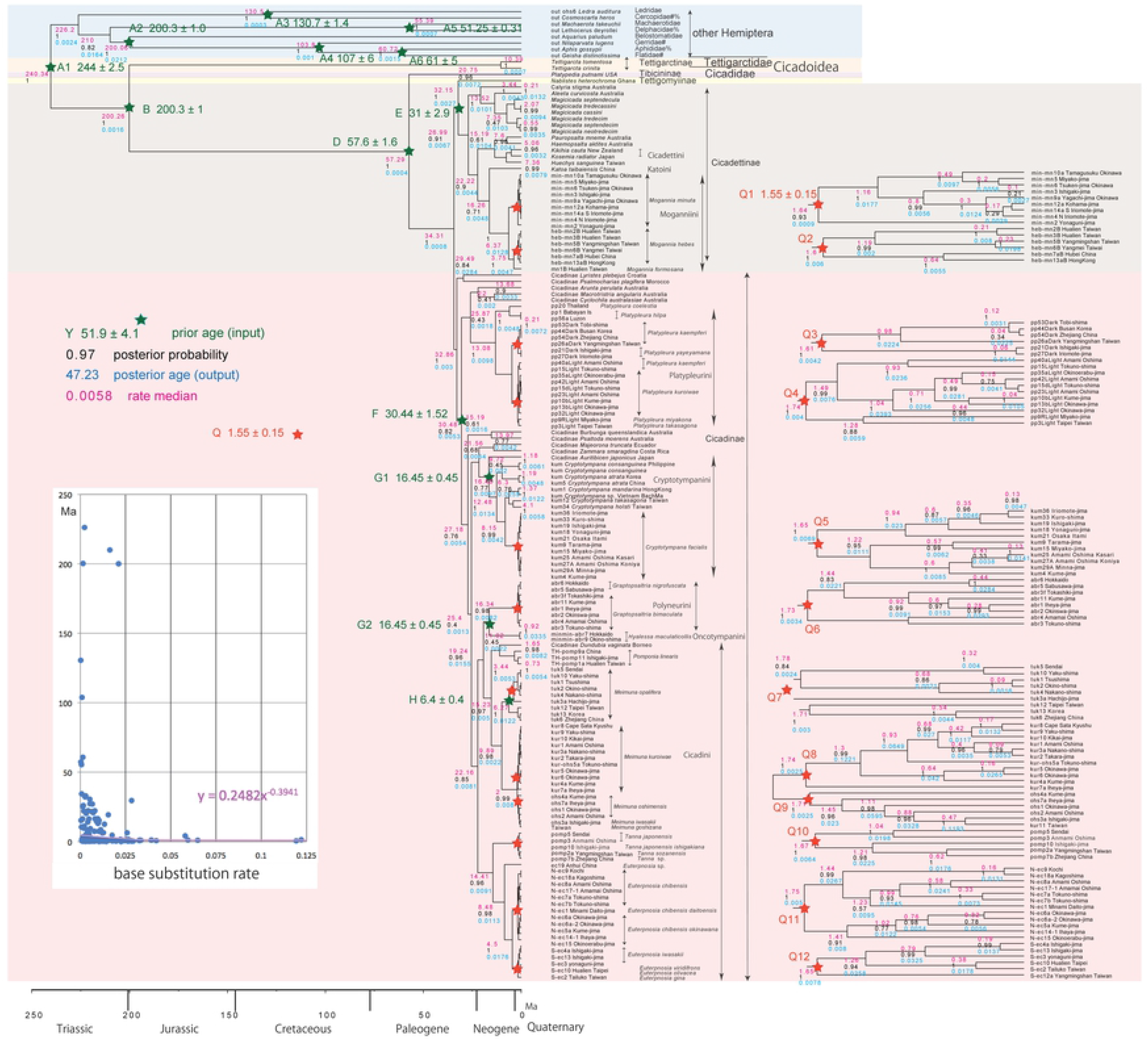
Cicada timetree built by BEAST v1.X, applying 1,534 bp COI and 874 bp 18S rRNA sequences. OUTs with isolate number: our own analyzed specimens shown in Table 1, and others: from GenBank / DDJB. In outgroup Hemiptera, #: analyzed family by Johnson et al. (2018), % analyzed family by Misof et al. (2014). Inserted figure: Base substitution rate (= rate median shown at each node; mutations per bp per million years) vs age (= posterior age shown at each node) diagram. Purple curve with equation: Exponential trendline calculated using Excel.

Phylogenetic trees by Marshall et al. (2018) and Łukasik et al. (2018) were not dated phylogenies, whereas the latest version of BEAST (Bayesian Evolutionary Analysis Sampling Trees; Suchard et al., 2018) v1.X released on 10th June 2018 has a fossil age calibration protocol. This calibration involves applying times of the most recent common ancestors (tMRCAs) of the ingroup species in the associated software of BEAUti (Bayesian Evolutionary Analysis Utility). Minimum and maximum age constraints in MCMCtree is complexly defined (Benton & Donoghue, 2007; Marshall, 2008), and application of the latter is not easy, but it is not needed in BEAST v.1.X. Moulds (2018), one of authors of Marshall et al. (2018), reviewed the ages of cicada fossils, and these dates up to 200 Ma were available for our fossil-based time calibrations in BEAUti. As shown by Osozawa et al. (2017), *Platypleura* and some other cicadas can be additionally and simultaneously calibrated by the Quaternary geological event calibration at 1.55 Ma. In addition, partitioning in BEAUti is performed simply by applying each gene sequence, not by concatenated sequences found in all the previous studies including Marshall et al. (2018). Using this most recent version of BEAST (v1.X), we can build an accurate and reliable cicada timetree.

Through these analyses, we corroborate the classification and some rearrangement of species into four subfamilies of Tibicininae, Tettigomyiinae, Cicadettinae, and Cicadinae included in a family of Cicadidae by Marshall et al. (2018) and Łukasik et al. (2018), and then estimate the crown dates of these subfamilies and tribes, and the dates of every species (Fig. 1). In the present analyses, we include *Derotettix*, a relict species of new subfamily of Derotettiginae with the oldest lineage in Cicadidae (Simon et al. 2019), and also try to estimate the crown date. Comparison in the context of the whole Hemipteroid insect timetree (Johnson et al., 2018) and whole insect timetree (Misof et al. 2014) can be conducted as an extension of this analysis, adding other Hemiptera species as outgroup.

Our main purpose was to represent precise evolutionary history of whole cicadas by contracting the BEAST timetree, but we have additional bellow because we have new attractive finding. Recent phylogenetic analyses usually employs relaxed clock model, which allows each branch of a phylogenetic tree to have its own evolutionary rate (Drummond et al. 2012). Although the relaxed distribution can be set such as lognormal in BEAUti, the rate of variability has not been considered till the present study. Because the output figure of BEAST v.1X represents the base substitution rate and age at each node, we can check the acceleration or delay of rate through the time on the newly built timetree, as the first trial. Chronogram by Marshall et al. (2016) for Australian Cicadettini might be unsuccessful, probably because of unsuitable calibration method and software choice.

In our previous experience of *Platypleura* cicadas calibrated solely by 1.55 Ma (Osozawa et al. 2017), our outgroup choice of *Tettigarcta crinita* output the extremely young origin as old as 10 m.y. We can now explain that the one digit slow base substitution rate of old time than the present affected on the apparently young date of the *Tettigarcta* lineage.

We show that the increasing cicada diversity reflects exponentially increasing base substitution rate approaching recent times (Figs. 1 ~ 3 inset), that may be a consequence of extensive adaptive radiation triggered by the start of the Quaternary glacial-interglacial cycles and severe environmental and climatic change. Although Tettigarctidae is a living fossil as old as 200 Ma, our work shows that the Cicadinae subfamily differentiation mostly took place after 40 Ma, and the speciation is relatively younger event. The latter is partly a consequence of the Quaternary vicariant speciation of East Asian cicada in the Ryukyu and Taiwan region (e.g., Osozawa et al. 2017), but rather expresses the increased biodiversity reflecting the recently accelerated base substitution rate and speciation including the cryptic species found in each island or island group.

## Materials and Methods

### Taxon sampling and applied DNA sequence

COI and 18S rRNA sequence data from our collected 113 specimens are in Table 1. Primers used, amplifications, and sequencing are given in Osozawa et al. (2017). These sequences were aligned by ClustalW in MEGA X (Kumar et al. 2016). The COI sequence data comprise 1,534 bp, and the 18S rRNA sequence 874 bp, and the resolution to construct phylogenetic tree is enough, as we experienced in the previous *Platypleura* paper (Osozawa et al. 2017). The COI data in Marshall et al. (2018) comprised 1485 pb comparable to ours, and their COII data comprised 684 pb. Nuclear 18S rRNA represented less variation with much slower base substitution rate compared to mitochondrial COI, and the topology was unaffected (Osozawa et al. 2017). North American Cryptotympanini was analyzed by Hill et al. (2015), applying 1467 bp of COI and 783 bp of nuclear EF-1a, and the resolution was enough. Mostly Australian Cicadettini was analyzed by Marshall et al. (2016), applying 1492 bp of COI and 1047 bp of nuclear EF-1a, and the resolution was also enough.

We incorporated other representative cicada data from GenBank / DDJB, derived from Marshall et al. (2018) and Łukasik et al. (2019), and included these data in our BEAST analyses. Note that Tettigarctinae, Derotettiginae, Tibicininae, and Tettigomyiinae are not known from East Asia, and Cicadettinae has only two species of *Kosemia* in Japan main islands. By this restriction, our own worldwide taxon sampling was very difficult, and GenBank / DDJB data offered by Marshall et al. (2018) and Łukasik et al. (2019) were essential for us. Whole mitochondrial sequence data by Łukasik et al. (2019) are included in our analyses as corresponding COI regions. However, note that not all data found in Marshall et al. (2018) corresponded to our COI and 18S rRNA sequences, and these data were not available in our analyses. Within COI sequence data in Marshall et al. (2018), only 21 data for Cicadettinae and only 14 data for Cicadinae are available in our analyses even by considering data found in Łukasik et al. (2019). 18S rRNA sequence in Marshall et al. (2018) is only available for *Nablistes heterochroma* of Tettigomyiinae and *Platypedia putnami* of Tibicininae. Marshall et al. (2018) additionally offered sequence data of COII, ARD1, and EF-1a, but these data in their table 1 were not enough to cover all their OTUs on figure 4. In the first step, we made a timetree only applying COI sequence data (Figs. 1 and 2; 191 OTUs) that covers Tettigomyiinae and Tibicininae species. Next, we made a timetree generated by applying both COI and 18S rRNA sequences (Fig. 3; 155 OTUs). We confirmed that our analyses with or without 18S rRNA data did not affect on the topologies.

In our analyses, single OUT did not represent single species (Table 1). *Platypleura* cicada (Osozawa et al. 2017) was experienced severe vicariance affected by the 1.55 isolation of the Ryukyu, Japan, and Taiwan islands from Chinese continent (Osozawa et al. 2012), and we needed to collect each island population for each *Platypleura* species. Similarly, we newly collected cicada specimens in this paper form each island population of *Mogannia* (Cicadettinae), and *Cryptotympana*, *Graptopsaltria*, *Hyalessa*, *Pomponia*, *Meimuna*, *Tanna*, and *Euterpnosia* (Cicadinae). *Hyalessa maculaticollis* was affected by vicariance within China and Japan (Liu et al. 2018). As noted above, Tettigarctinae, Derotettiginae, Tibicininae, and Tettigomyiinae are lacking in East Asia, and we could not collect these cicadas.

### Phylogenetic analyses associated with fossil and geological event calibrations by BEAST v1.X

A Bayesian inference (BI) tree (Figs. 1 ~ 3) was constructed using the software BEAST v.1X, running BEAUti, BEAST, TreeAnnotator, and FigTree, in ascending order. Before operating the BEAST software, the BEAGLE Library must be downloaded. Tracer v.1.6 was applied for checking the calculation status and estimating such as mean base substitution rate.

For graphic explanation of the operation of this software, see the “BEAST operating manual” at: http://kawaosombgi.livedoor.blog/archives/11386037.html

In BEAUti, “Taxa” and “Priors” (date setting is in “Prior”; date should be set in box of “tMRCA”) were set as follows for ingroup species relative to two partitions of COI and 18S rRNA. These partitions automatically appear in “Partitions” without traditional PartitionFinder, if these sequence data are uploaded by using the plus button or by “Import Data”.

Calibrations points are shown on Figs. 1 and 2, and these dates were input in “Priors” in BEAUti; they are summarized below. Corresponding ingroup species were included in ingroup taxa by “Taxon Set” on the “Taxa” screen in BEAUti. Other than calibration points A1 to A6 for the outgroup (Fig. 1), the fossil calibrations are after Moulds (2018). For the latter, some are based on radio-isotopic dating the fossil-bearing strata, whereas others are based on biostratigraphy assigned to an age/stage on the geologic time scale, for which absolute age ranges are generally based on radio-isotopic dates of associated strata in key global localities. This time scale has been standardized by the International Commission on Stratigraphy (ICS) (www.stratigraphy.org) and the most recent version of the time scale is available at http://www.stratigraphy.org/index.php/ics-chart-timescale, and the explanatory paper related to the generation of the time scale is Cohen *et al*. (2013). Calibration points Q1 to Q12 are after our geological event calibration that adopts a 1.55 Ma date based on multiple biostratigraphic and radio-isotopic dates connected to various geologic relationships in the Ryukyu islands region (Osozawa *et al*. 2012): this geologic event calibration was used in a previous study of *Platypleura* cicadas by our group (Osozawa et al. 2017).

The shape of the underlying chronogram was controlled and affected by the calibration points and dates in the BEAUti setting as shown by prior age (input) in Figs. 1 ~ 3, and in this meaning the BEAST calibration was “active” one. If the shape was unreasonable including a case of output age was distinct from input age, we reset the calibration and repeatedly run BEAST until to get reasonable tree. We show below the final calibration points and their dates.

The specific calibration points and dates used are as follows:

Calibration point A1: Crown Hemiptera: Fossil Aphidoidea was reported from the French Bundsandstein (Szwedo and Nel 2010; Bashkuev et al. 2012) of Anisian age (244.5 ± 2.5 Ma).

A2: The oldest fossil Belostomatidae was reported from the Zagaje Formation, Poland (Popov 1996) of Hettangian age (200.3 ± 1.0 Ma).

A3: Fossil Cicadellidae including Ledridae (Zhang 1997) and fossil Cercopidae (Hong 1982) was from the Jehol Biota, northern China. The Jehol Biota horizon has been dated by the Ar-Ar method on associated silicic tuff at 130.7 ± 1.4 Ma (He et al. 2006).

A4: Fossil Gerridae is from French amber (Perrichot et al. 2005) of Albian age (107 ± 6 Ma).

A5: Fossil Delphacidae is from the Green River Formation, USA (Grande 1980). Ar-Ar dating on silicic tuff within the formation yields ages of 53.5 – 48.5 Ma (weighted average age of 51.25 ± 0.31 Ma; Smith et al. 2003).

A6: Fossil Flatidae was from the Maíz Gordo Formation, northwest Argentina (Petrulevičius 2011) of Paleocene age (61 ± 5) Ma.

Calibration point B: Crown Tettigarctinae: Fossil Tettigarctinae was found in strata Dorset, England (Whalle 1985) of Hettangian age (203.1± 1.0 Ma).

Calibration point C: Stem Derotettiginae: Food plant of *Derotettix mendosensis* is Amaranthaceae in Argentina (Simon et al., 2019), and this worldwide C4 plant was phylogenetically studied by Piirainen et al. (2017). This fossil was reported by Zucol et al. (2018), and the fossil horizon was dated by Ar-Ar method at 49.512 ± 0.019 Ma (Eocene; Woodburne et al., 2014). However, fossil *Burmacicada protera* was found from Burmese amber (Poinar and Kritsky 2011), and the zircon U-Pb age of the amber bearing matrix was 98.79 ± 0.62 Ma (Shi et al. 2012). We applied this date for stem Cicadidae and stem Derotettiginae.

Calibration point D: Stem *Platypedia putnami* (Tibicininae): Fossil *Platypedia primigenia* was found in the Florissant Formation, Colorado, USA, and associated strata was dated by Ar-Ar method at 35.15 ± 1.65 Ma (Mcintosh et al. 1992). However, we used older date of crown Tibicininae: Fossil *Davispia bearcreekensis* was found in the Upper Fort Union Formation, Montana, USA (Cooper 1941), and the age was considered the Thanetian of 57.6 ± 1.6 Ma (Flores and Bader 1999).

Crown Cryptotympanini: Fossil *Hadoa grandiose* was also found in the Florissant Formation, Colorado, USA, and associated strata was dated by Ar-Ar method at 35.15 ± 1.65 Ma (Mcintosh et al. 1992), but this calibration generated unreasonable tree.

Calibration point E: Crown Cicadettinae: *Paracicadetta oligocenica* (Boulard and Nel 1990) was from Cereste, France, and this famous fossil locality was considered to be the Ruperian of 31 ± 2.9 Ma.

Calibration point F: Stem *Lyristes plebejus*: Fossil Lyristes sp. was reported from Seifhennersdorf, Germany, and associated strata was dated by the K-Ar method as 30.44 ± 1.52 Ma (Walther and Kvacek 2007).

Calibration point G: Crown *Cryptotympana*: Fossil *Cryptotympana incasa* and *C. miocenica* (G1), and also *Hyalessa lapidescens* (G2) were found from Shanwang, Shandong, China, and these strata are considered to be time correlative to the European MN5 mammalian unit (16.45 ± 0.45 Ma; Roček et al. 2011).

Calibration point H: Crown *Meimuna opalifera*: Fossil *Meimuna protopalifera* was found from the Itamuro Formation, Tochigi, Japan (Fujiyama 1982; Yoshikawa 2005), and the fission track age of the correlative terrestrial strata of the Nashino Formation, Sendai, is 6.4 ± 0.4 Ma (Fujiwara et al. 2008).

Calibration point Q: The dates of the common ancestors of *Mogannia minuta* (Q1), *M. hebes* (Q2), *Cryptotympana facialis* (Q3), dark winged *Platypleura* (Q4), right winged *Platypleura* (Q5; Osozawa et al. 2017), *Graptopsaltria nigrofuscata* + *G. bimaculata* (Q6), *Meimuna opalifera* (Q7), *Meimuna kuroiwae* (Q8)*, M. oshimaensis + M. iwasakii* + *M. goshizana* (Q9), *Tanna japonensis* + *T. japonensis ishigakiana* + *T. sozanensis* + *T.* sp. (Q10) *Euterpnosia chibensis* + *E. chibensis daitoensis* + *E. chibensis okinawana* (Q11), *E. iwasakii* + *E. viridifrons* + *E. olivacea* + *E. gina* + *E.* sp. (Q12): The date assigned is that of a geological event, which is of isolation of the Ryukyu islands from the Chinese continent by the opening of the Okinawa trough that began (i.e., islands had separated from mainland and each other by this time) at 1.55 ± 0.15 Ma (Osozawa et al. 2012). The age assignment is from multiple biostratigraphic and radio-isotopic ages from the oldest marine strata on the landward side of the islands as well as the sides facing other islands, so that the age of such strata constrain the physical separation of the islands from the mainland and each other. Note that no land bridge for dispersal was geologically evidenced in the Ryukyu islands.

## Results

### Hemiptera Timetree (Fig. 1)

Because topology is concordant between Figs. 1 and 2 (COI) and Fig. 3 (COI + 16S rRNA), we describe bellow based on Fig. 2 with 191 OTUs, although each posterior age (output) was not all the same. Based on our analyses, the subfamily-group classification follows Marshall *et al*. (2018), Łukasik et al. (2019), and Simon et al. (2019).

Hemiptera, including Cicadoidea, has a single common ancestor of 243.11 Ma, as calibrated by the 244.5 ± 2.5 Ma age reviewed above. Phylogeny of the outgroup Hemiptera calibrated by A1 to A6 was concordant to Johnson et al. (2018) and Misof et al. (2014).

In the Cicadoidea ingroup, Tettigarctidae was an old lineage differentiated from Cicadidae at 200.25 Ma, as calibrated by 200.3 ± 1 Ma, so Tettigarctidae is essentially a living fossil that have persisted since ca. 200 Ma. We estimated a date of the common ancestor of two extant species of *Tettigarcta tomentosa* (Tasmania) and *T. crinita* (southeast Australia) at 15.05 Ma, and the youngest fossil of Tettigarctinae was reported from the Aquitanian (21.735 ± 1.295 Ma), southern New Zealand (Kaulfuss & Moulds, 2015). However, Tettigarctidae includes 19 extinct genera according to Kaulfuss1 and Moulds (2015), whereas modified as many more according to Moulds (2018).

Simon et al. (2919) proposed a new subfamily Derotettiginae consisting of a single species of *Derotettix mendosensis*, which is a sister of resting Cicadidae species and the oldest lineage species in Cicadidae dated at 98.85 Ma. Łukasik et al. (2018) already showed such the phylogenetic position of *D. mendosensis*.

Our timetree showed that Tibicininae is a sister of Tettigomyiinae + Cicadettinae + Cicadinae differentiated at 65.89 Ma, and Tibicininae started differentiation at 57.07 Ma. Tettigomyiinae is a sister of Cicadettinae differentiated at 35.67 Ma, Tettigomyiinae + Cicadettinae is a sister of Cicadinae differentiated at 39.44 Ma. Cicadettinae started differentiation at 31.48 Ma, Cicadinae started differentiation at 36.7 Ma. Differentiation of Tettigomyiinae + Cicadettinae took place simultaneously after 39.44 Ma.

A single common ancestor of Cicadidae excepting Derotettiginae started differentiation and speciation into Tibicininae, Tettigomyiinae, Cicadettinae, and Cicadinae at 65.89 Ma. Although the pre-Miocene fossil Cicadidae collectively include ten extinct genera, comprising *Davispia* and *Lithocicada* for Tibicininae, *Paracicadetta*, *Paleopsalta*, *Minyscapheus*, and *Miocenoprasia* for Cicadettinae, and *Burmacicada, Camuracicada*, *Tymocicada*, *Dominicicada* for Cicadinae, the remaining 23 genera post-Oligocene fossil cicadas are extant (Moulds 2018). Cicadidae, conflict to the fossil record, consisted of only one species but coexisted with a Tettigarctidae species between 200.25 and 65.89 Ma, and cicada biodiversity was extremely low during this period excepting for extinct species and *D. mendosensis*.

The geologic calibration point Q1 to Q12 at 1.55 ± 0.15 Ma applies to multi furcations that were recognized for *Mogannia minuta* and other cicadas endemic to in the Ryukyu islands and Taiwan as noted above. Each island or island group population was mostly genetically distinct, endemic, and criptic, as shown for *Platypleura* in Osozawa et al. (2017).

### Inconsistent cicada base substitution rate (Figs. 1 ~3 insets)

Comparing base substitution rate vs age shows that the rate has not been constant; the rate appears to have exponentially increased approaching the present. The data points, fitted trendline, and associated equation are shown on the insets of Figs. 1 and 2. The trendlines and associated rates are similar for analyses based on COI alone (Figs. 1 and 2), and combined COI + 18S rRNA (Fig. 3).

## Discussion

### Recently increased cicada biodiversity

Hemipteroid insects of Psocodea, Thysanoptera, and the subject of this study, Hemiptera, include 120,000 described species which comprise over 10% of known insect diversity, date back to 400 Ma (Hemiptera: 300 Ma; Johnson et al. 2018). Johnson et al. (2018) estimated primarily Cretaceous dates of differentiation into species, including Cercopoidea, Gerriidea, Flatidae, and Cicadoidea, that are common to our analyses, although they analyzed only two to nine superfamily taxa, in contrast to the 180 taxa of Cicadoidea and 8 taxa for other Hemiptera of our analyses. Misof et al. (2014) estimated mostly in pre-Paleogene dates of differentiation into species including Cercopoidea, Aphididae, and Delphacidae common to our analyses, with less than 13 taxa analyzed. Their higher level phylogeny suggested long branches and old lineage of each super family species concordant to ours, and did not suggest the geologically recent increase in insect diversity apparent from our analyses of 191 Hemiptera taxa.

In Fig. 2, ingroup Cicadoidea, excluding Derotettiginae, underwent extensive differentiation into 180 taxa after 65.89 Ma, mostly after 39.44 Ma, leading to increasing biodiversity of Cicadoidea. Cicadoidea consisted of only two species excluding *D. mendosensis* between 98.85 and 65.89 Ma, although Cicadoide contains many extinct species that remain to be identified as fossils (Moulds 2018).

Increased biodiversity was typically presented by criptic species in each island of Ryukyu. For example, *Platypleura kaempferi* in the Amami and Okinawa islands has light colored wings contrasting to that in Japan-Korea-China and Taiwan with dark colored wings, and the clade was distinct each other (Osozawa et al., 2017). *P. kaempferi* is not a single species but includes at least two criptic species of light or dark winged *Platypleura*. Cicadas calibrated by other Q points included cryptic species, which also contributed increasing biodiversity. The Okinawa trough is spreading now, the Ryukyu islands are separating from the Chinese continent, and the vicariantly speciation and radiation is under progress, which is also contributing increasing biodiversity. In China mainland, *Hyalessa maculaticollis* and *Platypleura hilpa* were extensively radiated to form cryptic species (Liu et al., 2018, 2020).

### Exponentially increased base substitution rate as a factor of Hemiptera diversity, and their possible causes

Figs. 1 ~ 3 insets show a large range of base substitution rates for different time periods, at variance with the constant molecular clock hypothesis (relatively constant rate over time; Ho 2008). The trend in base substitution rates shows an exponential increase approaching recent time.

Such an increase in base substitution rate was first shown for taxa such as primates by Ho et al. (2005) who showed that a Quaternary calibration date resulted in a more a rapid base substitution rate than that associated an older calibration date. They employed an older version of BEAST (v1. 3; Drummond and Rambaut, 2003) that required repeated runs, applying a date at each calibration point. In contrast, BEAST v1.X, used in our analyses, can simultaneously apply multiple calibration points, as we have done using dates ranging from the Triassic to the Quaternary. As a result, the calculated increasing rate of base substitution in our analyses is not an artifact of a Quaternary calibration, but is constrained by multiple age calibrations across a wide range of geologic time. Therefore, although the base substitution rate trendlines and associated equations of Ho et al. (2005) are similar to ours, their timetrees do not reflect the changing of base substitution rates through time, but rather reflect a constant base substitution rate as if constrained by a strict molecular clock. A similar situation relative to our BEAST analyses was for beetles in the Aegean region reported by Papadopoulo et al. (2010).

The increasing base substitution rate is apparently associated with the recently increasing cicada diversity, expansion, and radiation (in Fig. 2 timetree) that started at 39.44 Ma. The timing of the most rapid diversification coincides with Quaternary environmental change, marked by the start of glacial-inter glacial cycles. The initiation of Quaternary glaciations may have been triggered by rapid expansion of land grasses (Poales), that led to increased carbon fixation that decreased atmospheric CO2 concentrations, because of the high efficiency of CO2 fixation of such C4 plants (Taira, 2007; Sage, 2004). C4 Poales appeared and began diversification during the Oligocene (23 – 33.9 Ma) based on molecular clock approach, and after 14.5 Ma based on fossil evidences (Sage 2004). We estimated 15.58 Ma by our Angiospermae timetree, also employed BEAST v1.X with robust plant fossil calibrations (Osozawa et al., 2020).

Feeding plants of *D. mendosensis* are, however, C4 dicots of Amaranthaceae (see figure 9 in Sage 2004) and Chenopodiaceae in degraded salt-plain habitats in arid regions of central Argentina (Simon et al. 2019). These dicot fossils but including C4 monocot fossils of Poales grass (Chloridoidae) were reported from the Eocene in Patagonia by Zucol et al. (2018), **and** the fossil horizon was dated by the Ar-Ar method of 49.512 ± 0.019 Ma (Woodburne et al. 2014). The C4 photosynthetic pathway began from 50 Ma in South America, earlier than elsewhere. As for Chloridoideae, however, transition from C3 to C4 photosynthesis occurred in the Oligocene (23~33.9 Ma) was reported by Christin et al. (2008), but Sage (2004) suggested the fossil date 14.5 Ma similar to the above.

The ultimate factor of increasing biodiversity was thus generation and radiation of C4 plants and development of grass lands on the Earth since the Oligocene but probably middle Miocene, through the decreasing atmospheric CO2 concentrations, start of the Quaternary ice ages, adaptive radiations, and increasing base substitution rates. The Earth environment was thus much affected by life activity including C4 grasses, and if no life, the Earth might have become the completely different bleak planet.

## Supplementary Material

Supplementary data in Table 1 are available at GenBank / DDBJ.

## Data Availability Statement

All relevant data are within the manuscript.

## Acknowledgements

We thank Chris Simon and David Marshall for privately offering Tettigarcta sequence data (later released in GenBank / DDBJ). We thank the collectors shown in Table 1. Bor-ming Jahn (Taiwan University; deceased 1 December, 2016), Ping-Shih Yang (Taiwan University), Chin-Ho Tsai (National Dong Hwa University), and Jen-Zon Ho and Hua-Te Fang (Endemic Species Research Institute) supported sample collections and obtained permission to collect in Taiwan.

## Author Contributions

Soichi Osozawa collected samples and coordinated the research, carried out the DNA analyses, and wrote this paper, and John Wakabayashi (native English-speaking American geologist) edited the writing.

## References

Benton MJ, Donoghue PCJ. 2007. Paleontological evidence to date the tree of life. Molelular Biology and Evolution 24: 26–53.

Benton MJ, Donoghue PCJ, Asher RJ, Friedman MN, Thomas J, Jakob V. 2015. Constraints on the timescale of animal evolutionary history. Palaeontologia Electronica 18.1.1FC: 1–106.

Bashkuev A, Se J, Aristov D, Ponomarenko A, Sinitshenkova N, Mahler H. 2012. Insects from the Buntsandstein of Lower Franconia and Thuringia. Paläontologische Zeitschrift 86: 175–185.

Boulard M, Nel A. 1990. Sur deux cigales fossiles des terrains tertiaires de la France (Homoptera, Cicadoidea). Revue Francaise d’Entomologie 12: 37–45.

Cohen KM, Finney SC, Gibbard PL, Fan J-X. 2013. The ICS International Chronostratigraphic Chart. Episodes 36: 199–204.

Cooper KW. 1941. *Davispia bearcreekensis* Cooper, A new cicada from the Paleocene, with a brief review of the fossil Cicadidae. American Journal of Science 239: 286–304.

Drummond AJ, Suchard MA, Xie D, Rambaut A. 2012. Bayesian phylogenetics with BEAUti and the BEAST 1.7. Molecular Biology and Evolution 29: 1969–1973.

Flores RM, Bader LR. 1999. Fort Union coal in the Powder River Basin, Wyoming and Montana: A synthesis. U.S. Geological Survey Professional Paper 1625-A: 1–71.

Fujiyama I. 1969. A Miocene Cicada from Nasu, with an additional record of a Pleistocene cicada from Shiobara, Japan. Bulletin of the National Science Museum, Tokyo: 12: 863–875.

Fujiwara O, Yanagisawa Y, Irizuki T, Shimamoto M, Hayashi H. Danhara T, Fuse K, Iwano H. 2008. Chronological data for the Middle Miocene to Pliocene sequence around the southwestern Sendai Plain, with special reference to the uplift history of the Ou Backbone Range. Bulletin of Geological Survey Japan 59: 423–438.

Grande L. 1980. Paleontology of the Green River Formation with a review of the fish fauna. The Geological Survey of Wyoming Bulletin 63: 1–333.

He HY, Wang XL, Jin F, Zhou ZH, Wang F, Yang LK, Ding X, Boven A, Zhu RX. 2006. The 40Ar/39Ar dating of the early Jehol Biota from Fengning, Hebei Province, northern China. Geochemistry, Geophysics, Geosystems 7: Q04001. doi:10.1029/2005GC001083 ISSN: 1525-2027

Hill KBR, Marshall DC, Moulds MS, & Simon C. 2015. Molecular phylogenetics, diversification, and systematics of *Tibicen* Latreille, 1825 and allied cicadas of the tribe Cryptotympanini, with three new genera and emphasis on species from the USA and Canada (Hemiptera: Auchenorrhyncha: Cicadidae). Zootaxa 3985: 219–251.

Ho S. 2008. The molecular clock and estimating species divergence. Nature Education 1: 168.

Ho SY, Phillips MJ, Cooper A, Drummond AJ. 2005. Time dependency of molecular rate estimates and systematic overestimation of recent divergence times. Molecular Biology & Evolution 2: 1561–1568.

Hong YC. 1982. Mesozoic Fossil Insects of Jiuquan Basin in Gansu Province. Clapham. In Chinese.

Johnson KP et al. 2018. Phylogenomics and the evolution of hemipteroid insects. Proceedings of the National Academy of Sciences of the United States of America: 115: 12775–12780.

Kaulfuss U, Moulds M. 2015. A new genus and species of tettigarctid cicada from the early Miocene of New Zealand: *Paratettigarcta zealandica* (Hemiptera, Auchenorrhyncha, Tettigarctidae). ZooKeys 484: 83–94.

Liu Y, Qiu Y, Wang X, Yang H, Hayashi H, Wei C. 2018. Morphological variation, genetic differentiation and phylogeography of the East Asia cicada *Hyalessa maculaticollis* (Hemiptera: Cicadidae). Systematic Entomology 43: 308–329.

Liu Y, Pham HT, He Z, Wei C. 2020. Phylogeography of the cicada *Platypleura hilpa* in subtropical and tropical East Asia based on mitochondrial and nuclear genes and microsatellite markers. International Journal of Biological Macromolecules 151: 529–544.

Łukasik P, Nazario K, Van Leuven JT, Campbell MA, Meyer M, Michalik A, Pessacq P, Simon C, Veloso C, McCutcheon JP. 2018. Multiple origins of interdependent endosymbiotic complexes in a genus of cicadas. Proceedings of the National Academy of Sciences of the United States of America 115: E226–E235.

Marshall DC. 2008. A simple method for bracketing absolute divergence times on molecular phylogenies using multiple fossil calibration points. American Naturalists 171: 726–42.

Marshall DC, Hill KBR, Moulds MS, Vanderpoo, D, Cooley JR, Mohagan AB. Simon C. 2016. Inflation of molecular clock rates and dates: Molecular phylogenetics, biogeography, and diversification of a global cicada radiation from Australasia (Hemiptera: Cicadidae: Cicadettini). Systematic Biology 65: 16–34.

Marshall DC, Moulds M, Hill KBR, Price BW, Wade EJ, Owen CL, Goemans G, Marathe K, Sarkar V, Cooley JR, Sanborn AF, Kunte K, Villet MH, Simon C. 2018. A molecular phylogeny of the cicadas (Hemiptera: Cicadidae) with a review of tribe and 926 subfamily classification. Zootaxa 4424: 1–64.

Misof B et al. 2014. Phylogenomics resolves the timing and pattern of insect evolution. Science 346: 763–767.

Mcintosh WC, Geissman JW, Chapin CE, Kunk MJ, Henry CD. 1992. Calibration of the latest Eocene-Oligocene geomagnetic polarity time scale using 40Ar/39Ar dated ignimbrites. Geology 20: 459–463.

Moulds MS. 2018. Cicada fossils (Cicadoidea: Tettigarctidae and Cicadidae) with a review of the named fossilised Cicadidae. Zootaxa 4438: 443–470.

Osozawa S, Shinjo R, Armig R, Watanabe Y, Horiguchi T, Wakabayashi J. 2012. Palaeogeographic reconstruction of the 1.55 Ma synchronous isolation of the Ryukyu Islands, Japan, and Taiwan and inflow of the Kuroshio warm current. International Geology Review 54: 1369–1388.

Osozawa S, Shiyake S, Fukuda H, Wakabayashi J. 2017. Quaternary vicariance of Platypleura (Cicadidae) in Japan, Ryukyu, and Taiwan islands. Biological Journal of the Linnean Society 121: 185–199.

Papadopoulo A, Anastasiou I, Vogler AP. 2010. Revisiting the insect mitochondrial molecular clock: The mid-Aegean trench calibration. Molecular Biology and Evolution 27: 1659–1672.

Perrichot V, Nel A, Néraudeau D. 2005. Gerromorphan bugs in Early Cretaceous French amber (Insecta: Heteroptera): first representatives of Gerridae and their phylogenetic and palaeoecological implications. Cretaceous Research 26: 793–800.

Petrulevičius JF. 2011. Paleogene insects from Maíz Gordo Formation, northwest Argentina: Taphonomy, diversity and paleobiogeography, 337–347. In Salfity JA, Marquillas RA, edts., Cenozoic Geology of the Central Andes abd Argentina SCS.

Piirainen M, Liebisch, O, Kadereit G. 2017. Phylogeny, biogeography, systematics and taxonomy of Salicornioideae (Amaranthaceae / Chenopodiaceae) – A cosmopolitan, highly specialized hygrohalophyte lineage dating back to the Oligocene. TAXON 66: 109–132

Poinar Jr G, Gene Kritsky G. 2011. Morphological conservatism in the foreleg structure of cicada hatchlings, *Burmacicada protera* n. gen., n. sp. in Burmese amber, *Dominicicada youngi* n. gen., n. sp. in Dominican amber and the extant *Magicicada septendecim* (L.) (Hemiptera: Cicadidae). Historical Biology 1: 1–6.

Popov YA. 1996. The first record of a fossil water bug from the Lower Jurassic of Poland (Heteroptera: Nepomoprha: Belostomatidae). Polskie Pismo Entomologiczne 65: 101–105.

Sage RF. 2004. The evolution of C4 photosynthesis. New Phytologist 161: 341–370.

Shi G, Grimaldi DA, Harlow GE, Wang J, Wang J, Yang M, Lei W, Li Q, Li X. 2012. Age constraint on Burmese amber based on U-Pb dating of zircons. Cretaceous Research 37: 155–163.

Simon C, Gordon ERL, Moulds MS, Cole JA, Haji D, Lemmon AR, Lemmon EM, Kortyna M, Nazario K, Wade EJ, Meister RC, Goemans G, Chiswell SM, Pessacq P, Veloso C, Mccutcheon JP, Łukasik P. 2019. Off-target capture data, endosymbiont genes and morphology reveal a relict lineage that is sister to all other singing cicadas. Biological Journal of the Linnean Society 128: XX–YY.

Smith ME, Singer B, Carroll A. 2003. 40 Ar/39 Ar geochronology of the Eocene Green River Formation, Wyoming. Geological Society of America Bulletin 115: 549–565.

Suchard MA, Lemey P, Baele G, Ayres DL, Drummond AJ, Rambaut A. 2018. Bayesian phylogenetic and phylodynamic data integration using BEAST 1.10. Virus Evolution 4: vey016.

Szwedo J, Nel A. 2011. The oldest Aphid insect from the Middle Triassic of the Vosges, France. Acta Palaeontologica Polonica 56: 757–766.

Taira A. 2007. Search the Earth History, Geology 3, pp. 396. Iwanami Shoten, Publishers, Tokyo. (in Japanese)

Walther H, Kvacek Z. 2007. Early Oligonene flora of Seifhennersdorf (Saxiny). Acta Musei Nationalis Pragae, Series B, Historia Naturalis 63: 85–174.

Whalley PES. 1985. The systematics and palaeogeography of the Lower Jurassic insects of Dorset, England. Bulletin of the British Museum of Natural History (Geology) 39: 107–189.

Woodburne MO, Goin FJ, Raigemborn MS, Heizler M, Gelfo JN, Oliveira EV. 2014. Revised timing of the South American early Paleogene land mammal ages. Journal of South American Earth Sciences 54: 109–119.

Yoshikawa T. 2005. Paleoenvironmental change and age of the Miocene Kanomatazawa Formation in the Shiobara area, Tochigi Prefecture, central Japan. Journal of Geological Society of Japan 111: 39–49. In Japanese with English abstract.

Zhang H. 1997. Early Cretaceous insects from the Dalazi Formation of the Zhixin Basin, Jilin Province, China. Palaeoworld 7: 75–103.

Zucol AF, Krause JM, Brea M, Raigemborn MS, Matheos SD. 2018. Emergence of grassy habitats during the greenhouse–icehouse systems transition in the Middle Eocene of southern South America. Ameghiniana 55: 451–482.

